# The natural axis of transmitter receptor distribution in the human cerebral cortex

**DOI:** 10.1101/2020.09.28.316646

**Authors:** Alexandros Goulas, Jean-Pierre Changeux, Konrad Wagstyl, Katrin Amunts, Nicola Palomero-Gallagher, Claus C Hilgetag

**Affiliations:** Institute of Computational Neuroscience, University Medical Center Hamburg-Eppendorf, Hamburg, Germany; Collège de France and Institut Pasteur CNRS, Paris, France; McGill Centre for Integrative Neuroscience, Montréal Neurological Institute, Montréal, Canada; Department of Psychiatry, University of Cambridge, Cambridge, United Kingdom, Wellcome Trust Centre for Neuroimaging, University College London, London, United Kingdom; Institute of Neuroscience and Medicine (INM-1), Research Centre Jülich, Jülich, Germany; C. and O. Vogt Institute for Brain Research, University Hospital Düsseldorf, Heinrich Heine University Düsseldorf, Düsseldorf, Germany; Department of Psychiatry, Psychotherapy, and Psychosomatics, Medical Faculty, RWTH Aachen, and JARA-Translational Brain Medicine, Aachen, Germany; Health Sciences Department, Boston University, 635 Commonwealth Ave. Boston, MA 02215, USA

**Keywords:** cortical organization, unifying principles, molecular diversity

## Abstract

Transmitter receptors constitute a key component of the molecular machinery for inter-cellular communication in the brain. Recent efforts have mapped the density of diverse transmitter receptors across the human cerebral cortex with an unprecedented level of detail. Here, we distil these observations into key organizational principles. We demonstrate that receptor densities form a natural axis in the human cerebral cortex, reflecting decreases in differentiation at the level of laminar organization, and a sensory-to-association axis at the functional level. Along this natural axis, key organizational principles are discerned: progressive molecular diversity (increase of the diversity of receptor density), excitation/inhibition (increase of the ratio of excitatory-to-inhibitory receptor density) and mirrored, orderly changes of the density of ionotropic and metabotropic receptors. The uncovered natural axis formed by the distribution of receptors aligns with the axis that is formed by other dimensions of cortical organization, such as the myelo- and cytoarchitectonic levels. Therefore, the uncovered natural axis constitutes a unifying organizational feature linking multiple dimensions of the cerebral cortex, thus bringing order to the heterogeneity of cortical organization.

---

**T**transmitter receptors are essential for cellular communication, since they are the molecular elements responsible for the responsiveness of cells to distinct types of neurotransmitters. Such key role has naturally attracted concerted efforts that aim to map and elucidate the role of receptors in a plethora of neurobiological phenomena, ranging from cellular to cognitive processes, with important ramifications for understanding pathologies involving disturbances at the receptor level and drug design (1–7). Positron emission tomography of the human brain can offer a coarse mapping of the distribution of certain receptors, useful for modeling receptor dynamics in the human brain (8) and understanding the pathogenesis of brain disorders at the molecular level (9). While densities for some receptors can be measured in humans with positron emission tomography, for instance, for serotonin (10), there are no suitable positron-emission tomography tracers that can map a plethora of key receptors in living humans in a safe way (e.g., for NMDA). Furthermore, the number of different receptor types that can be examined in a single individual is very small. In vitro quantitative receptor autoradiography of receptor densities has been used for a broad range of key transmitter receptors and offers spatial resolution suitable for uncovering details pertaining to the laminar architecture of the cerebral cortex (7, 11, 12). Moreover, receptor autoradiography reflects quantitative, neurobiologically interpretable measurements. Recent efforts have mapped the distribution of a broad range of transmitter receptors in multiple cortical areas across different laminar compartments (6, 7, 13). These advancements also pose the challenge of compressing these raw observations into a parsimonious set of comprehensible, key organizational principles.

Building on recent endeavors (7), here, we distill key organizational principles of the transmitter receptor distribution of the human cerebral cortex. We highlight an axis of maximum variance of the receptor distribution across the cortex. This natural axis ranges from sensory to association areas of the cerebral cortex. The arrangement of areas along this axis also entails three organizational principles. First, progressive molecular diversity is observed, that is, an increase of the diversity of receptor densities when proceeding from one end of the axis (sensory areas) to the other end of the axis (association areas). Second, progressive excitation/inhibition ratio, that is, the increase of the ratio of excitatory-to-inhibitory receptor density, is also observed along the same direction along the natural axis. Third, progressive, mirrored changes of the density of ionotropic and metabotropic receptors are observed, that is, the density of metabotropic receptors increases when proceeding from sensory to association areas, while the density of ionotropic receptors is mirrored, thus, decreasing along this direction. The aforementioned principles manifest in a laminar-wise fashion, showcasing the importance of uncovering organizational principles at a fine spatial resolution. Moreover, the uncovered receptor-based natural axis aligns with the spatially ordered changes of the cerebral cortex at other levels of architecture, such as the myelo- and cytoarchitectonic levels, in agreement with classic and more recent studies. Our results generate concrete testable predictions that are pertinent to cognitive, computational, and comparative neuroscience.

## Results

We analyzed the previously collected data of transmitter receptor densities across a wide range of areas of the human cerebral cortex (7). Density for multiple excitatory, inhibitory, ionotropic and metabotropic receptors has been measured (AMPA, NMDA, kainate, GABA_*A*_, GABA_*A*_/BZ, GABA_*B*_, M1, M2, M3, a_4_b_2_, a_1_, a_2_, 5-HT_1*A*_, 5-HT_2_, D_1_). Data were acquired in a laminar-wise fashion summarized in densities for granular, infra- and supragranular layers. For the receptor autoradiographical incubation protocols used and overall densitometric analytical procedure, see (7). An exemplary section is shown in Fig. 1 A.

**Fig. 1.**
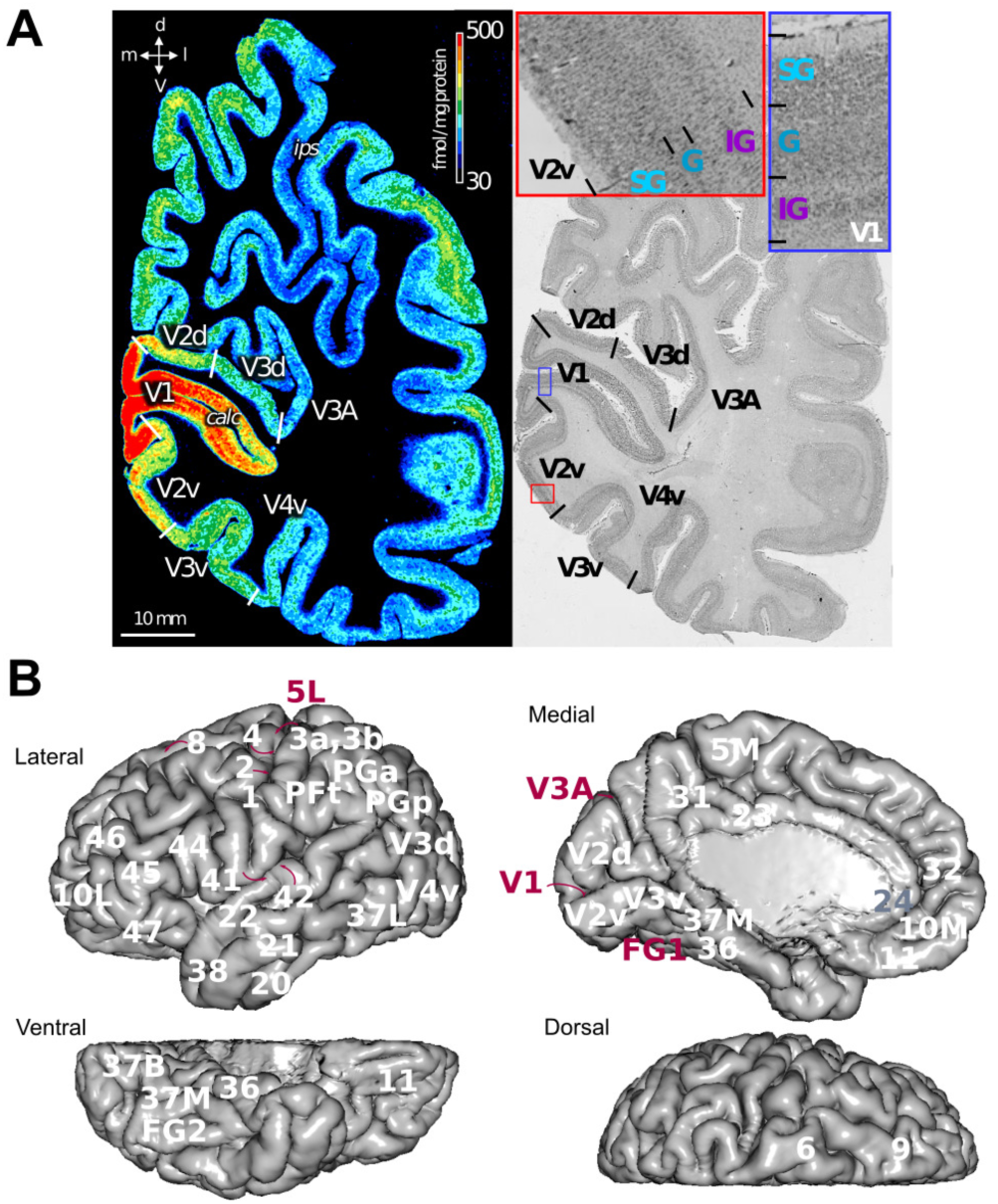
*In vitro* receptor autoradiography for estimation of transmitter receptor densities. A. Example of coronal sections through a human hemisphere, revealing the density of cholinergic muscarinic M2 receptors (left) and the distribution of cell bodies (right) for multiple visual areas. B. Summary of cortical areas for which laminar transmitter receptor measurements were carried out in (7, 11). Panel A adapted from (7, 11). SG=supragranular, G=granular, IG=infragranular.

### Uncovering the natural axis of transmitter receptor density

As a first step, we assembled the receptor density profile of each cortical area, that is, the density measured in each area for each of the aforementioned receptors in granular, infra- and supragranular layers (hereafter G, IG and SG respectively). Note that due to the agranular character (lack of the granular layer) of area 24, the only area with this feature in the set of the 44 cortical areas analyzed in (7) (Fig. 1 B), this area was not included in the analysis to have a consistent profile with IG, G, and SG measurements for all areas. Inclusion of area 24 and analysis based only on measurements on IG and SG layers led to qualitatively similar results. We aimed at uncovering the trajectory across which most variance of receptor density manifests. To this end, we applied Principal Component Analysis (PCA) to the receptor density profile of all cortical area. The arrangement of cortical areas in the space spanned by PC1 while PC2 unveiled a sensory-to-association axis (PC1, accounting for 28% of the variance) and PC2 was driven by the “outlier” character of areas 6 and 4, separating them from the rest of cortical areas (PC1, accounting for 22% of the variance) (Fig. 2). Note that here the terms “association” and “sensory” are not used to denote strict categories for grouping cortical areas. Instead, the terms are used to denote the broad character of areas that span the two extreme ends of the natural axis. In other words, the current results underline the natural axis as a continuum (or gradient) of receptor density variations and not as a discrete set of categories. The biplot simultaneously depicts the observations, that is, cortical areas (denoted with circles tagged with the names of each area), and the features, that is, laminar-wise receptor densities (arrows tagged with the names of each receptor and corresponding laminar compartment), in the PCA space (Fig. 2). Therefore, the biplot allows deciphering what receptor density measurements drive the segregation of cortical areas across the PCA space. Here, we will focus on PC1, which explains the largest amount of variance, and is, thus, referred to as the primary natural axis of receptor distribution.

**Fig. 2.**
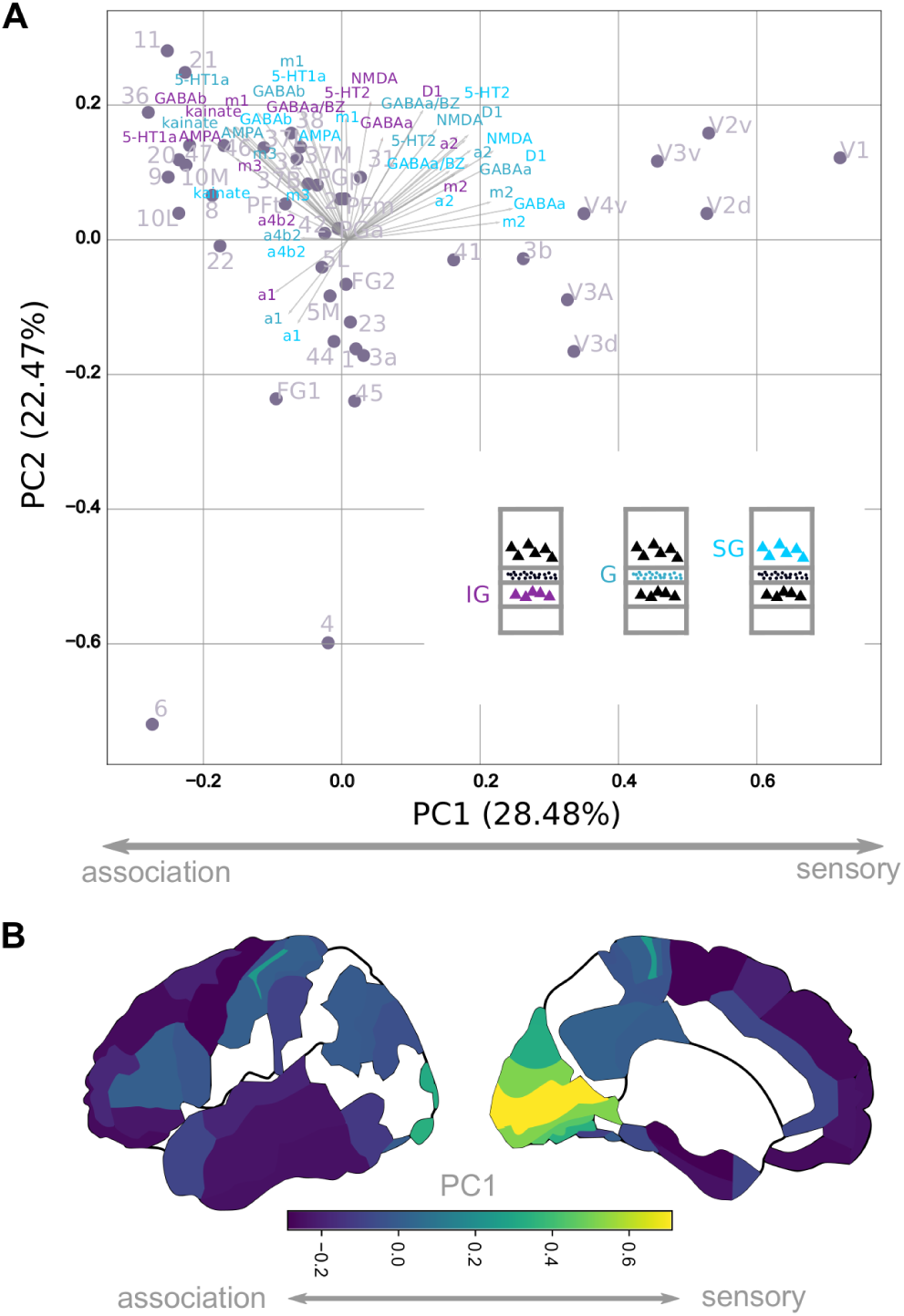
The natural axis of transmitter receptor density distribution. A. PC1 reveals the distribution of the areas along an axis explaining most of the receptor density variance across cortical areas. The axis stretches from sensory to association areas. We refer to this axis (PC1) as the primary natural axis of transmitter receptor density distribution. Note that here, the terms “association” and “sensory” denote the broad character of areas that span the two extreme ends of the natural axis. In this context, “association” refers to non-sensory areas. Note that different receptor densities across different laminar compartments, depicted as the PCA coefficients, have a more prominent role in the arrangement of different area constellations along the natural axis. For instance, receptor density of the IG layers is more prominent in association areas and NMDA in the primary areas, while AMPA is prominent in association areas. Lastly, a distinctive profile for primary areas is highlighted, that is, prominence of density of the GABA_*A*_, a_2_, M2 and D1 receptors. B. The natural axis rendered in a surface map of the human cortex. Colours correspond to the PC1 values. Note the “association” to “sensory” arrangement of areas along the natural axis.

Certain insights into the receptor signatures that segregate cortical areas along the primary natural axis can be discerned in the biplot (Fig. 2). First, receptor density of the IG layers is more prominent in segregating association areas from sensory areas along the natural axis (note the concentration of loadings related to IG measurements with magenta color in the left part of the biplot). Second, the biplot reveals a striking segregation along the natural axis with respect to the glutamate receptors AMPA and NMDA. NMDA is prominent on the sensory-related end of the natural axis, while AMPA is prominent in the association-related end. Third, the biplot reveals the distinctive receptor signatures of sensory areas, that is, the prominent role the GABA_*A*_, a_2_, m_2_ and D1 receptors in visual sensory areas from association areas (Fig. 2).

We subsequently aimed at distilling organizational principles that are not readily discernable from the biplot. Specifically, we estimated the diversity of receptor density of each cortical area as the Shannon entropy of the receptor profile of each area. Entropy was calculated as H = -sum(N.*log(N)/log(M), where N is the normalized receptor profile of the area with each entry denoting each receptor density value as a proportion to the overall receptor density of the area and M denoting the total number of features of the profile, that is, 45 (densities for 15 receptors for G, IG and SG layers) and log is the natural logarithm (hence, entropy was normalized to the [0 1] interval). We also estimated the excitation/inhibition ratio for each cortical area by dividing the sum of the density of all the excitatory receptors by the sum of the inhibitory receptors. Lastly, we estimated the changes of the receptor density across cortical areas for ionotropic and metabotropic receptors separately. For the assignment of receptors to the aforementioned categories, that is, excitatory, inhibitory, ionotropic and metabotropic, see (7). The aforementioned metrics were estimated on a laminar-wise basis. The insights provided by these analyses are detailed in the sections below.

### Progressive molecular diversity

We associated the entropy of the areas, that is, their receptor diversity, to the primary natural axis. This association was performed on a laminar-wise basis. The analysis reveals that the entropy of areas, is aligned with the natural axis. Specifically, for the IG layers the progression from sensory to association areas along the primary natural axis marked an increase of the entropy, and thus, diversity of receptors (rho=-0.79, p<0.01). This relation was less pronounced for the G layers (rho=-0.42, p<0.01) and statistically absent for the SG layers (rho=-0.13, p>0.1) (Fig. 3 A). A laminar-wise rank ordering of cortical areas based on their receptor entropy is provided in SI Appendix Fig. S1. These results indicate that the transition from sensory to association areas along the natural axis is accompanied by an increase of the diversity of the receptor profile that an area exhibits. This relation was more prominent in IG layers. We denote this phenomenon as progressive molecular diversity across the primary natural axis of the cerebral cortex. We also summarized the entropy of the receptor density across all areas on a laminar-wise basis. The highest entropy was observed for the IG layers, followed by the entropy in the G and SG layers. Statistically significant differences concerned the entropy of the IG receptors when compared to the entropy of the G and SG layers (two-sample Kolmogorov-Smirnov test, 0.45, 0.50, respectively, p<0.01) (Fig. 3 C).

**Fig. 3.**
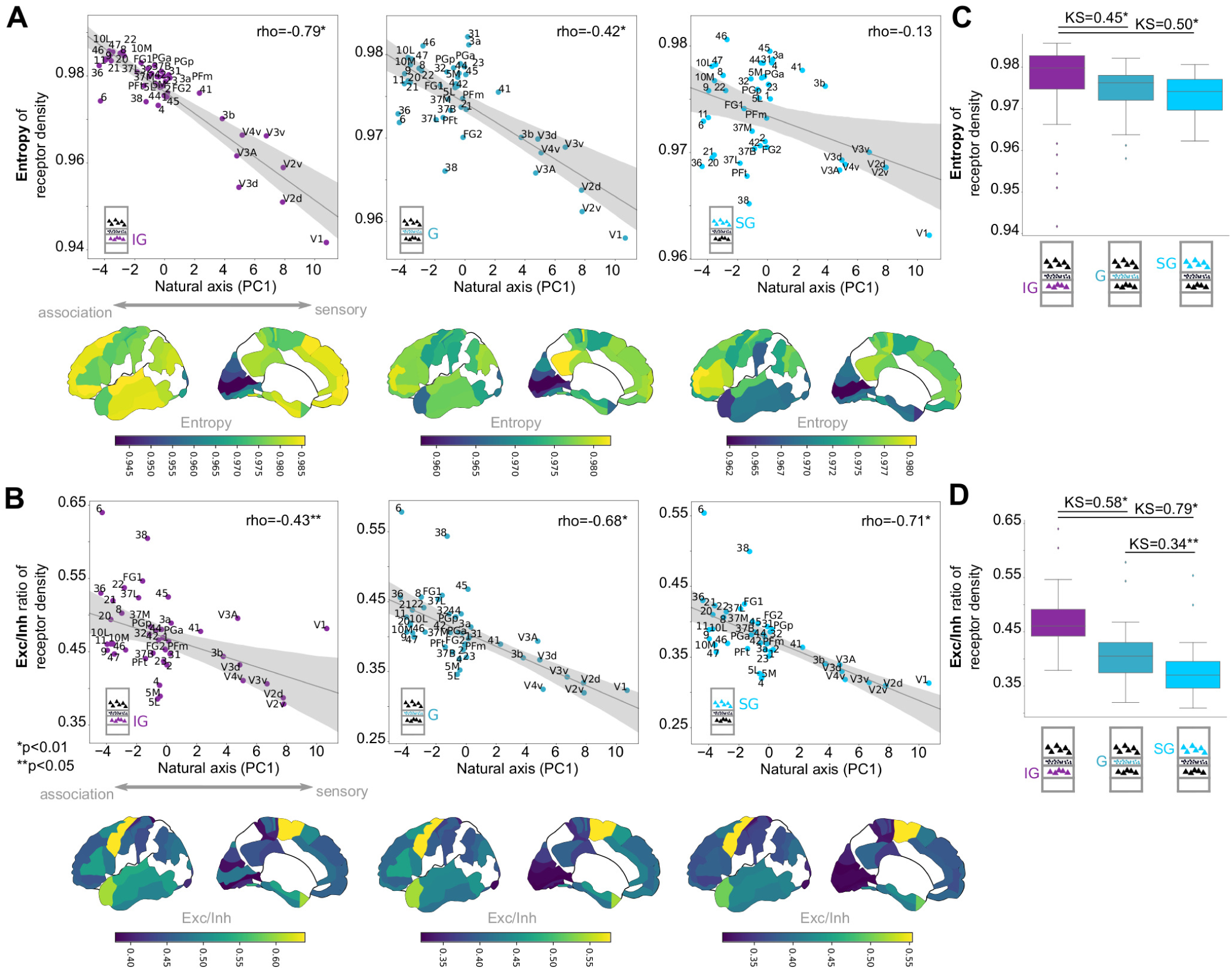
The principle of progressive molecular diversity and excitatory/inhibitory receptors ratio along the natural axis. A. Diversity of receptor profiles varies along the natural axis for each laminar compartment (IG, G, SG). Progressive molecular diversity refers to the increase of the entropy, thus diversity, of the receptor profile of areas when transitioning from sensory to association areas, especially prominent in IG layers. Bottom, surface maps depict the entropy values across the cerebral cortex. Note the sensory-to-association increase of entropy. B. Same as in (A), but for the excitatory/inhibitory receptors ratio. the ratio of excitatory/inhibitory receptors increases when transitioning from sensory to association areas, more prominently in SG layers. Note the sensory-to-association increase of the excitatory/inhibitory receptors ratio. C. Entropy of receptor profiles summarized for each laminar compartment (IG, G, SG) for all cortical areas. D. Same as in (C) but for the excitatory/inhibitory receptors ratio. Note that both the overall entropy and excitatory/inhibitory receptors ratio, summarized across all areas, is decreasing across IG, G, SG layers (panels C and D).

### Progressive molecular excitation/inhibition

We associated the excitation/inhibition of cortical areas to the primary natural axis. This association was performed on a laminar-wise basis. For all layers, the ratio of excitatory to inhibitory receptor density increased along the natural axis when transitioning from sensory to association areas (rho=-0.49, p<0.05, rho=-0.68, −0.71, p<0.01, for the IG, G and SG layers respectively) (Fig. 3 B). A laminar-wise rank ordering of cortical areas based on their excitation/inhibition ratio is provided in SI Appendix Fig. S2. Therefore, the transition from primary to association areas across the natural axis is accompanied by a progressive increase of the excitability of cortical areas. This progressive excitability was more prominent in SG layers. We denote this phenomenon as progressive molecular excitation/inhibition across the primary natural axis of the cerebral cortex. We also summarized the ratio of excitation/inhibition across all areas on a laminar-wise basis. The highest excitation/inhibition ratio was observed for the IG layers, followed by the excitation/inhibition ratio in the G and SG layers. Statistically significant differences concerned the excitation/inhibition ratio of the IG layers when compared to the entropy of the G and SG layers (two-sample Kolmogorov-Smirnov test, 0.58, 0.78, respectively, p<0.01), as well as the excitation/inhibition ratio of the G layers when compared to the entropy of the SG layers (two-sample Kolmogorov-Smirnov test, 0.34, p<0.05) (Fig.3 D).

### Mirrored density changes of ionotropic and metabotropic transmitter receptors

We next examined the relation of the density of ionotropic and metabotropic receptors to the primary natural axis. This analysis revealed mirrored changes of the density of the ionotropic and metabotropic receptors along the natural axis. Specifically, ionotropic receptor density increased when transitioning from association to sensory areas with the more prominent, statistically significant increase observed for the SG layers (rho=0.67, p<0.001). Contrary to the G and SG layers, the density of metabotropic receptors in IG decreased along the natural axis (rho=-0.49, p<0.001) (Fig. 4 A). The mirrored pattern was observed for the metabotropic receptors for all layers along the natural axis. Specifically, metabotropic receptor density decreased when transitioning from association to sensory areas, with the more prominent decrease observed for the IG layers (rho=-0.82, rho=-0.65, p<0.001, rho=-0.44, p<0.05, for the IG, G and SG layers respectively) (Fig. 4 B). A laminar-wise rank ordering of cortical areas based on the density of ionotropic and metabotropic receptors is provided in SI Appendix Fig. S3. In sum, the density of ionotropic receptors, on average, increases when transitioning from association to sensory areas, while the density of metabotropic receptors, on average, decreases. We denote this phenomenon as mirrored density changes of ionotropic and metabotropic receptors along the primary natural axis of the cerebral cortex.

**Fig. 4.**
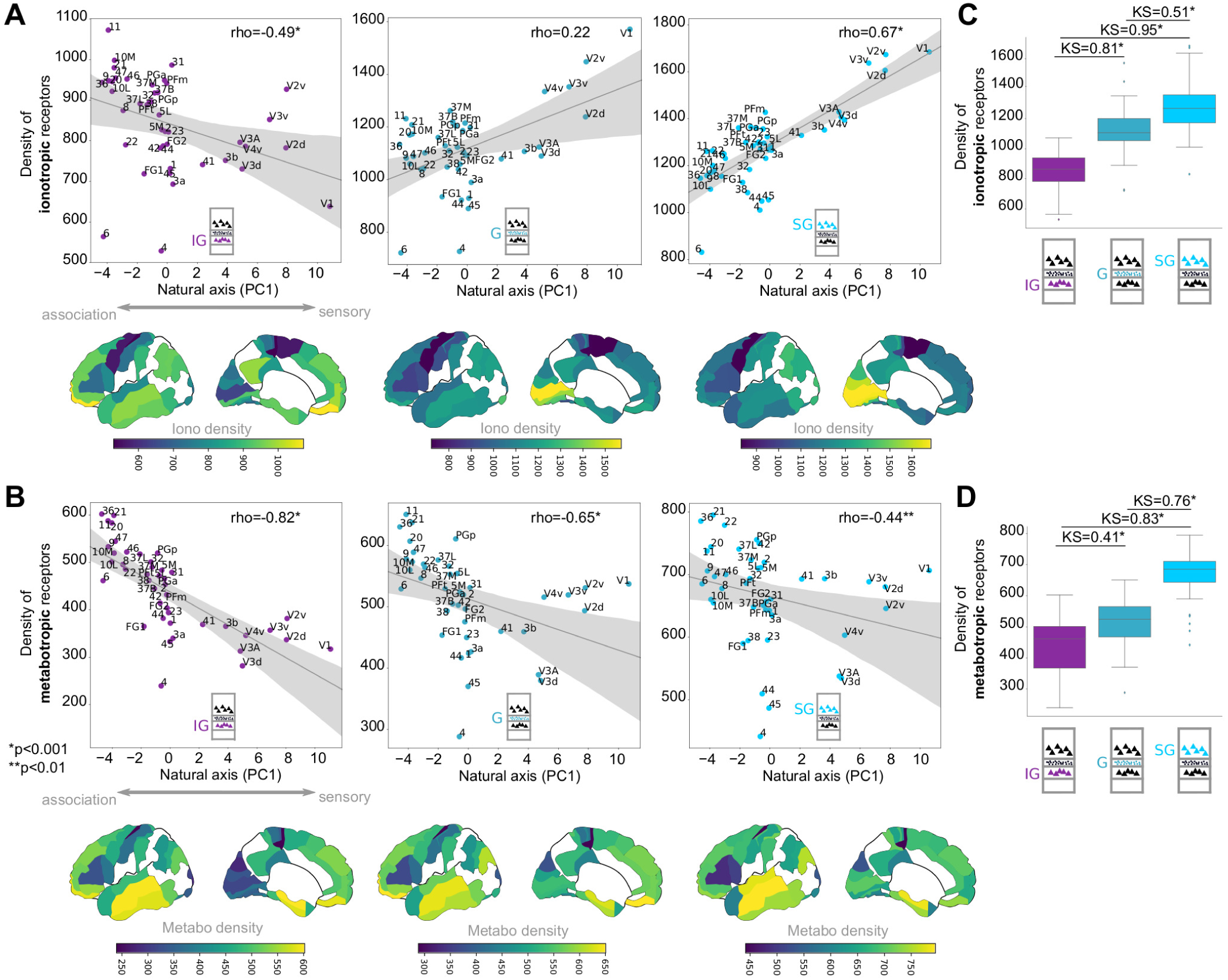
Mirrored density changes of ionotropic versus metabotropic transmitter receptors along the natural axis. A. Density of ionotropic receptors increases in the SG layers when transitioning from sensory to association areas. B. Same for (A) but for the metabotropic receptors. Note that the metabotropic receptor density exhibits a mirrored pattern along the natural axis when compared to the density of the ionotropic receptors (with the exception of IG layers). Note that there is an increase of metabotropic receptor density when transitioning from sensory to association areas, more prominently in IG layers, whereas there is a decrease of ionotropic receptor density when transitioning from sensory to association areas, more prominently in SG layers. C. Density of ionotropic receptors across all areas summarized for each laminar compartment (IG, G, SG). D. Same as in (C) but for the metabotropic receptors.

We also estimated the overall density for ionotropic and metabotropic receptors across cortical areas on a laminar-wise basis. Ionotropic receptors form transmembrane ligand-gated ion channels. Metabotropic receptors do not form channels, and their impact on the channels that a cell exhibits relies on signal transduction, that is, a cascade of intracellular events. Hence, on average, the effect of ionotropic receptors on the channels of the cell that they inhabit is faster when compared to the effect of metabotropic receptors. For both metabotropic and ionotropic receptors, an increase in the density was observed across IG, G and SG. These laminar-wise differences were statistically significant (two-sample Kolmogorov-Smirnov test Ionotropic: 0.81, 0.95, 0.51, Metabotropic: 0.41, 0.83, 0.76, for IG versus G, IG versus SG and G versus SG, respectively, p<0.001) (Fig. 4 C, D).

## Discussion

### The natural axis of transmitter receptor distribution

Numerous classic (14–18) and more recent studies (19–26) dictate that the heterogeneity of the cerebral cortex of mammals in general, and humans in particular, is not spatially random, but instead manifests as spatially ordered changes (*gradation principle* in (15)). These changes have a characteristic spatial trajectory with sensory areas at one end and association areas at the other end (14, 17, 27). This sensory to association axis also entails more to less laminar differentiation, that is, distinguishability and prominence of the layers of the cerebral cortex, for instance, in terms of cell packing density and/or size of cell bodies (14, 22, 26). Our results highlight that the distribution of transmitter receptor density also follows this spatial trajectory, and thus, forms a natural axis of variation at the molecular level. Therefore, this spatial trajectory at the molecular level is tightly linked with the spatial trajectory that pertains to the myeloarchitectonic (19), cytoarchitectonic (22) and functional (25) dimensions of cortical organization. These associations are yet another manifestation of the principle of concurrent change that pertains to the cerebral cortex, that is, variations of features across the cortical sheet are concurrent and involve multiple dimensions of cortical organization (27).

In particular, our analysis unveils the prominent receptors that drive the segregation of cortical areas along the natural axis. Specifically, glutamate receptors AMPA and NMDA exhibit opposed preferences: NMDA is prominent in the sensory, highly differentiated cortical areas, while AMPA is more prominent in less differentiated cortical areas. Moreover, these opposed preferences manifest in a layer-wise fashion. While the more pronounced NMDA preference towards highly differentiated sensory cortical areas involves the SG layers, the more pronounced AMPA preference towards less differentiated association cortical areas involves the IG layers. In other words, the laminar-wise relative preferences of AMPA and NMDA receptors reflect the shifts of the laminar-wise relative preference of termination of inter-areal connections, as empirical observations in the monkey and cross-species wiring principles dictate (20, 29–31) (Fig. 5 B). Such organizational principles substantiate prior suggestions in the context of the modeling of the global neuronal workspace hypothesis (32). These suggestions postulate that top-down connections (association to sensory) are more related to the propensity of NMDA receptors, whereas bottom-up connections (sensory to association) are more related to the propensity of AMPA receptors (32). Moreover, our results highlight the prominence of the GABA_*A*_, a_2_, M2 and D_1_ receptors for the highly differentiated sensory cortical areas. In sum, we reveal the natural axis of transmitter receptor distribution in the human cerebral cortex and the associated contributions of specific receptor types to the segregation of cortical areas along this axis.

**Fig. 5.**
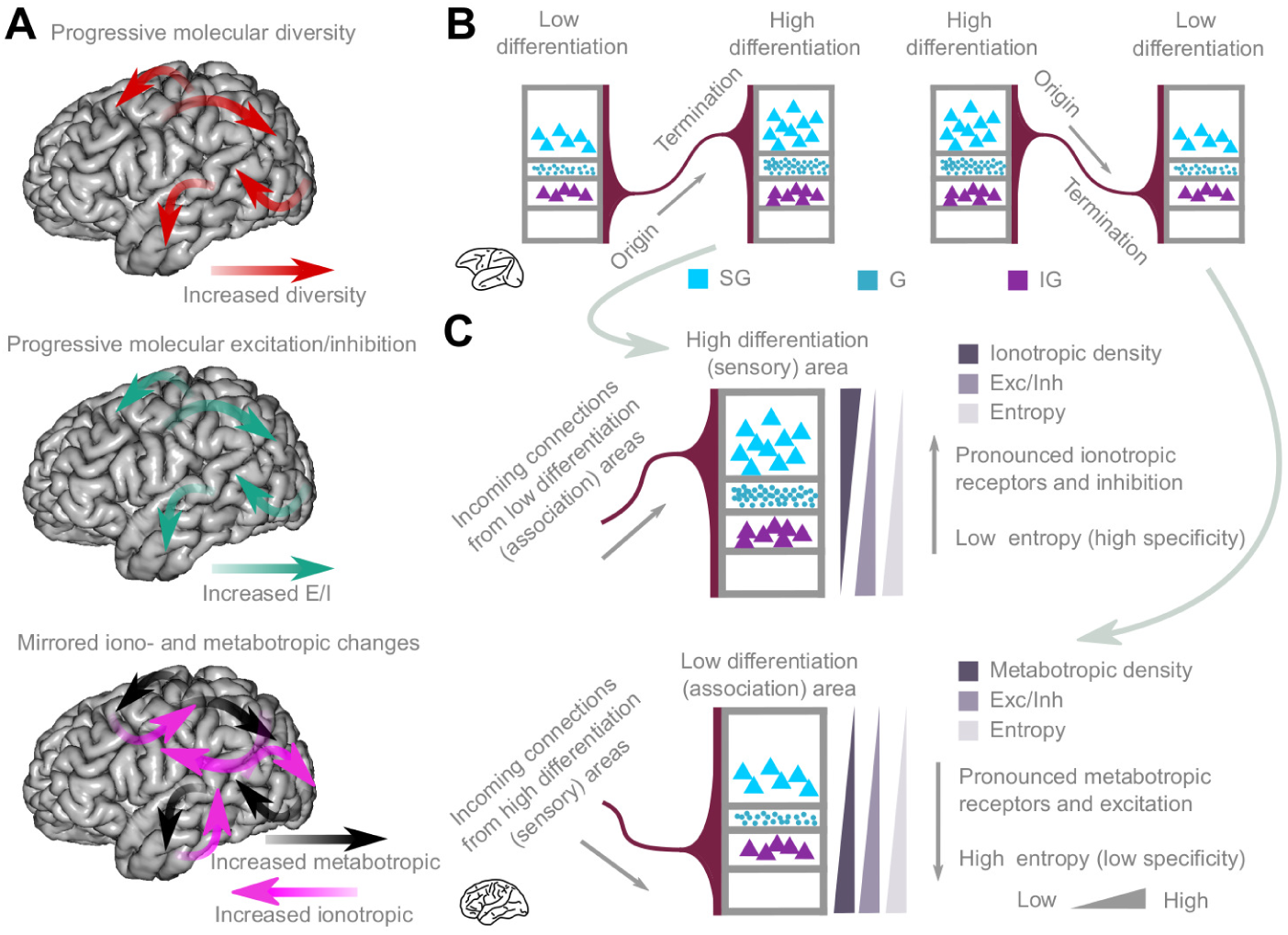
Organizational principles of receptor distribution and the inter-areal cortical projection system. A. Summary of organizational principles of receptor distribution along the primary natural axis. Note that this is a schematic representation denoting the overall changes across the sensory to association axis. For detailed results, see (Fig. 2, 3, 4). B. Wiring principles dictating the predominant origin and termination of cortico-cortical connections in the monkey. Less differentiated areas elicit connections predominantly from deep laminar compartments (IG layers) and when targeting more differentiated areas they terminate predominantly in upper layers (SG layers). The reverse pattern of origin and termination of connections is observed for efferents from more to less differentiated areas. The differentiation of areas is the most consistent explanatory model of the shifts of the laminar connectional patterns (28–30) and can be used for monkey to human extrapolations (20, 29). C. Receptor principles across layers for more and less differentiated areas. In more differentiated areas, towards the sensory end of the natural axis, the density of ionotropic receptors increases from deep (IG) to upper (SG) layers. Moreover, entropy and excitatory/inhibitory receptors ratio decreases, thus, a high specificity and an inhibitory nature prevails. These graded changes at the receptor level are aligned with the predominant terminations of connections from less differentiated areas to upper layers, as predicted by the monkey-based model in panel (B). In less differentiated areas, towards the association-related end of the natural axis, the density of metabotropic receptors increases from upper (SG) to lower (IG) layers. Moreover, entropy and excitatory/inhibitory receptors ratio increases, thus, a high molecular diversity and an excitatory nature prevails. These graded changes at the receptor level are aligned with the predominant terminations of connections from more differentiated areas to deep layers, as predicted by the monkey-based model in panel (B). Thus, the uncovered receptor principles distil insights that can be combined with predictions pertaining to the connectional level to unveil a molecular-connectional synergy in the human cerebral cortex.

### Progressive molecular complexification, excitability and mirrored ionotropic/metabotropic changes along the natural axis

We have highlighted three key organizational principles of the molecular composition of the cerebral cortex that manifest along the natural axis (Fig. 5 A). The principle of progressive molecular diversity illustrates that the diversity of receptor densities in each area increases when we transition from more differentiated, sensory areas to the less differentiated, association areas. This increase is more pronounced in IG layers. In other words, the entropy of the receptor profiles of each area increases along the natural axis. This principle indicates that association areas may exhibit their characteristic diverse functional profile, that is, their engagement in diverse tasks (33, 34), due to their broad tuning at the molecular level that allows them to be modulated by multiple neurotransmitter systems. Therefore, this principle offers cognitive neuroscience insights at the molecular level and generates a concrete prediction, that is, entropy of receptor profiles of cortical areas will be predictive of their functional diversity, above and beyond factors such as connectomic properties of areas (34). Moreover, combining insights from invasive tract-tracing studies in the monkey, cross-species wiring principles (20, 29–31) we can also predict that the predominant laminar termination of inter-areal structural connections will involve the IG layers of the human cerebral cortex and the current principle of molecular diversity (Fig. 5 C). This termination preference mirrors the increased entropy of the molecular profile of IG layers in association areas (Fig. 5 C). Therefore, we have a synergy between the connectional and receptor levels of organization that bestow the IG layers of association areas with a prominent role in accommodating incoming signals from areas that lie towards the other end of the natural axis. On the contrary, the preferential termination in more differentiated, sensory areas involves the SG layers (Fig. 5 B). SG layers of these areas exhibit low entropy, thus, their molecular profile is specialized and not broad (Fig. 5 C). In sum, the principle of progressive molecular diversity unifies receptor and connectional features and provides insights into the functional profile of cortical areas at the molecular level.

The principle of progressive excitation/inhibition ratio denotes that the excitation/inhibition ratio of receptor densities increases when we transition from more differentiated, sensory areas to the less differentiated, association areas. Observations in human and non-human primates indicate that the spine density of pyramidal cells increases in the transition from sensory to association areas (35). This feature bestows association areas with higher degrees of excitability that may contribute to their integrative functional capacity, as empirical and computational studies indicate (36–38). The principle of progressive increase in the excitatory/inhibitory receptors ratio extends these observations by offering a receptor-based explanation for the pronounced excitability of the neuronal populations inhabiting association areas. Such a principle can be instantiated in computational models embodying regional heterogeneity (36, 39, 40), thus offering enhanced neurobiological interpretability. Notably, the highest excitatory/inhibitory receptors ratio is observed in IG layers (Fig. 5 C). In monkeys, the predominant termination of inter-areal connections in association areas involves the IG layers (30) (Fig. 5 B). Inter-areal connections, especially connections spanning long distances are very weak in terms of the number of axons that they involve (31). Therefore, the highest excitability of IG may constitute a molecular signature that facilitates these weak, long-range connections to excite their downstream targets in the human cerebral cortex. Advancements in the *in vivo* mapping of long-range connections in humans may confirm such connectional-molecular synergy in the human cerebral cortex. Further predictions based on the principle of progressive molecular excitation/inhibition pertain to the functional connectivity properties of cortical areas. In monkeys and humans, prior studies involving a small set of cortical areas highlight a relation between the excitation/inhibition ratio and the functional connectivity strength of areas (41, 42). Thus, the principle of progressive excitation/inhibition ratio constitutes a template for predicting such functional connectivity properties of areas at a global whole-cortex level. From a clinical standpoint, the principle of progressive excitation/inhibition ratio across the cortical sheet may offer explanations and predictions with respect to pathology. For instance, areas with high excitation/inhibition ratio, such as areas 6 and 38, may be situated close to an excitability threshold above which pathological neuronal activity may arise, as is the case in epilepsy. In sum, the principle of progressive excitatory/inhibitory ratio can be instantiated in computational models to offer enhanced neurobiological interpretability and may predict and explain deviant neurophysiological profiles in a clinical context.

Lastly, the mirrored ionotropic/metabotropic changes along the natural axis dictate that the density of ionotropic receptors increases when we transition from association-to-sensory areas, while density of metabotropic receptors exhibits the mirrored pattern, that is, increases when we transition from sensory-to-association areas. Primary areas operate on fast temporal scales and the predominant density of ionotropic transmitter receptors may bestow them with this functional property, since the very nature of ionotropic receptors is the presence of ion channels with fast, immediate effects on the cell that they inhabit. On the contrary, association areas are characterized by high density of metabotropic transmitter receptors that have a slower, indirect impact on the cells that they inhabit. This property may bestow these areas with a functional repertoire that operates at slow temporal scales (such as learning processes) possibly accounting for their integrative and coordinating role. Such suggestion is further supported by empirical studies reporting higher concentrations of the enzyme calcium calmodulin-dependent protein kinase II in association areas of the monkey cortex (43). Notably, the increase of metabotropic receptor density towards the end of the natural axis occupied by association areas is more prominent in the IG layers, while the increase of ionotropic receptor density towards the other end of the natural axis occupied by sensory areas is more prominent in the SG layers (Fig. 5 A,C). Based on observations in the monkey and cross-species wiring principles (20, 29), for primary and association areas, these changes at the molecular level are centered around the most prominent laminar terminations, and thus, incoming signals, of inter-areal cortical connections (Fig. 5).

### Conclusions

We uncovered organizational principles that pertain to the distribution of the density of transmitter receptors in the human cerebral cortex. We demonstrate that receptor densities are organized along a natural axis in the cerebral cortex, ranging from sensory to association areas. The constellation of areas along this axis is characterized by organizational changes that are regulated by the principles of progressive molecular diversity, excitation/inhibition and mirrored changes of the density of ionotropic and metabotropic receptors. Our results showcase the importance and feasibility of taming the complexity of the cerebral cortex by distilling key organizational principles that bring order to the heterogeneity of cortical organization.

## Materials and Methods

Receptor autoradiography data that were used for this study were collected as described in (7). For analyses and metrics used, see SI Appendix SI Text. Python code used for the analysis and figures is available at: https://github.com/AlGoulas/receptor_principles

## ACKNOWLEDGMENTS

Funding from the European Union’s Horizon 2020 Research and Innovation Programme, grant agreement no. 604102 (HBP, SGA1) (KA), No. 785907 (HBP SGA2) (KA, CCH, JPC) and No. 945539 (HBP SGA3) (KA), as well as funding from the Deutsche Forschungsgemeinschaft (SFB 936/A1; TRR 169/A2; SPP 2041/HI 1286/7-1, HI 1286/6-1) (CCH) is gratefully acknowledged.

