## Supplementary material for "The natural axis of transmitter receptor distribution in the human cerebral cortex": SI Appendix SI Text

### Supporting Information Text

**Materials.** Receptor autoradiography data that were used for this study were collected as described in (1). Briefly, cortical areas were defined based on the Juelich-Duesseldorf atlas in combination with areas from Brodmann's map (2). The process involves the following steps. Autoradiographs, obtained from the protocol previously described (1), were digitized by means of a dedicated image acquisition and processing system. The relationship between the gray value of a pixel in the digitized autoradiograph and the receptor binding site density was defined by the gray value images of the co-exposed scales and were used to compute a calibration curve by non-linear, least-squares fitting, thus, defining the relationship between gray values in the autoradiographs and concentrations of radioactivity. Subsequently, these concentrations of radioactivity were corrected to account for experimental conditions, e.g., specific activity, dissociation constant and free concentration of the ligand during incubation. Thus, the gray value of each pixel in a digitized autoradiograph, which can be color coded for visualization purposes, codes for a receptor concentration per unit protein at saturation of ligand-receptor complexes (1). In total, data for 15 receptors, including, ionotropic, metabotropic, excitatory and inhibitory receptors, were obtained (AMPA, NMDA, kainate, GABA$_A$, GABA$_A$/BZ, GABA$_B$, M1, M2, M3, a$_4$b$_2$, a$_1$, a$_2$, 5-HT$_{1A}$, 5-HT$_2$, D$_1$) (1).

**Methods.** For each cortical area, a receptor profile was assembled that corresponds to the quantitative autoradiography measures for infragranular (IG), granular (G) and supragranular (SG) layers. We aimed at treating areas as observations and the laminar-wise measurements as features, to gain an understanding of the what features drive the areas to form an axis dictated by the highest variance in the receptor profile of the areas. Cingulate area 24 is agranular, therefore, it was excluded for the analysis. Qualitatively similar results were obtained when including areas 24 and using measurements only for IG and SG layers.

To uncover the axis of maximum variance, we performed a principal component analysis (PCA) on the z-scored features, that is, densities of receptors. To gain insights in a visually comprehensive fashion, a biplot that simultaneously depicts the scores and coefficients from the PCA analysis. PC1 was the focus of the report and is refereed to as the primary natural axis of receptor distribution. A tailored analysis aimed at examining principles that are not disclosed by PCA. Specifically, we hypothesized that areas might differ in the the degree of molecular diversity, that is diversity of receptor profiles. Previous studies have uncovered that functional diversity of association areas, in relation to primary areas, is related to connectional diversity (3). Therefore, we aimed at unravelling a molecular diversity property that differentiates association from primary areas. To this end, we computed the entropy of the receptor profiles of each area. Entropy for each area was calculated as H = -sum(N.*log(N)/log(M), where N is the normalized receptor profile of the area with each entry denoting each receptor density value as a proportion to the overall receptor density of the area and M denoting the total number of features of the profile, that is, 45 (densities for 15 receptors for G, IG and SG layers) and log is the natural logarithm (hence, entropy was normalized to the [0 1] interval, with 1 denoting the highest diversity). We also estimated the excitation/inhibition ratio for each cortical area by dividing the sum of the density of all the excitatory receptors by the sum of the inhibitory receptors. Previous studies have uncovered variations of the excitation/inhibition ratio in monkeys and humans and associated functional ramifications (4, 5). Moreover, excitation/inhibition balance is a prominent component of computational studies, and organizational principles based on excitation/inhibition are valuable for more neurobiologically informed and interpretable computational models of the human cortex. Lastly, excitation/inhibition balance imbalances are associated to pathological cases (6). Therefore, we aimed to uncover an organizational principle centered on the excitation/inhibition of the cerebral cortex. Lastly, we estimated the changes of the receptor density across cortical areas for ionotropic and metabotropic receptors separately. Due to the immediate, fast and indirect slower effect of ionotropic and metabotropic receptors, respectively, we hypothesized that a gradient of receptor density will be uncovered, with primary areas exhibiting the highest ionotropic densities, whereas the association would be characterized by high metabotropic receptor densities. For the assignment of receptors to the aforementioned categories, that is, excitatory, inhibitory, ionotropic and metabotropic, see (1).

The aforementioned metrics were estimated on a laminar-wise basis. All the aforementioned measurements were associated with the primary natural axis of receptor distribution (PC1). This association was estimated with Spearman's rank correlation coefficient. All analyses were performed with code written in Python.

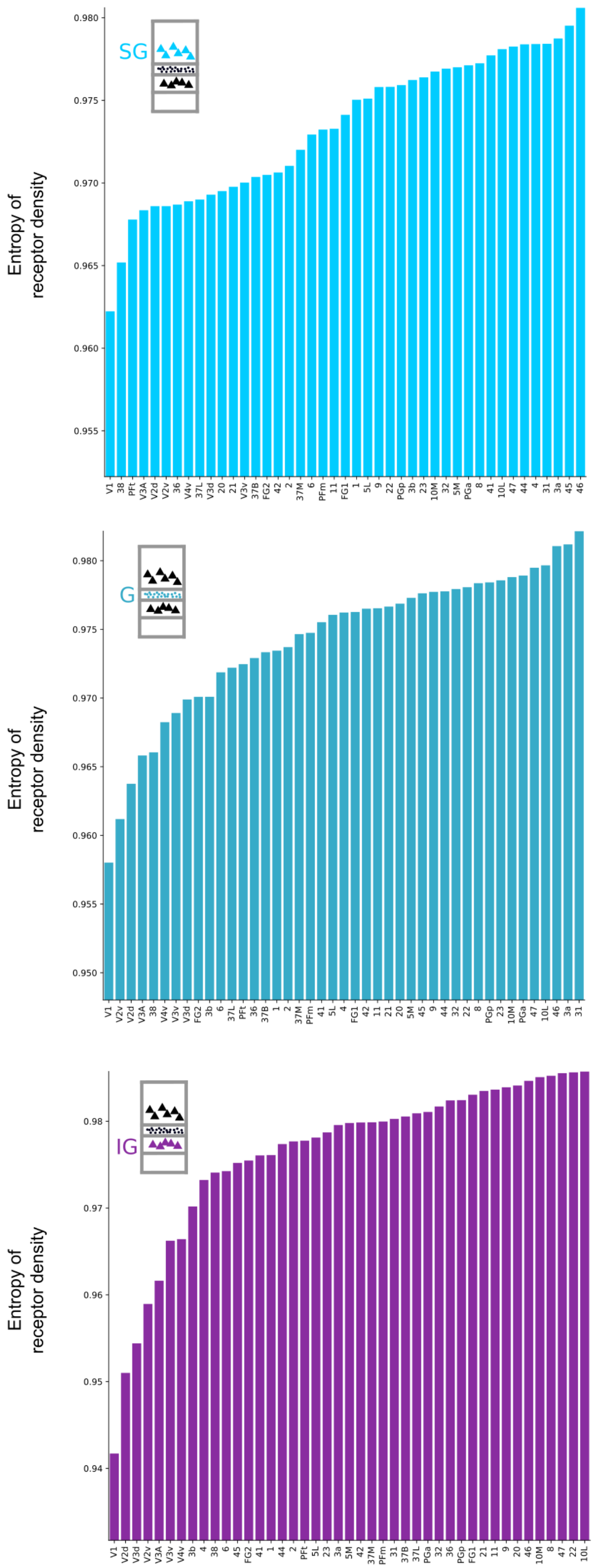


**Fig. S1.** Rank ordered areas based on their entropy in IG, G and SG layers.

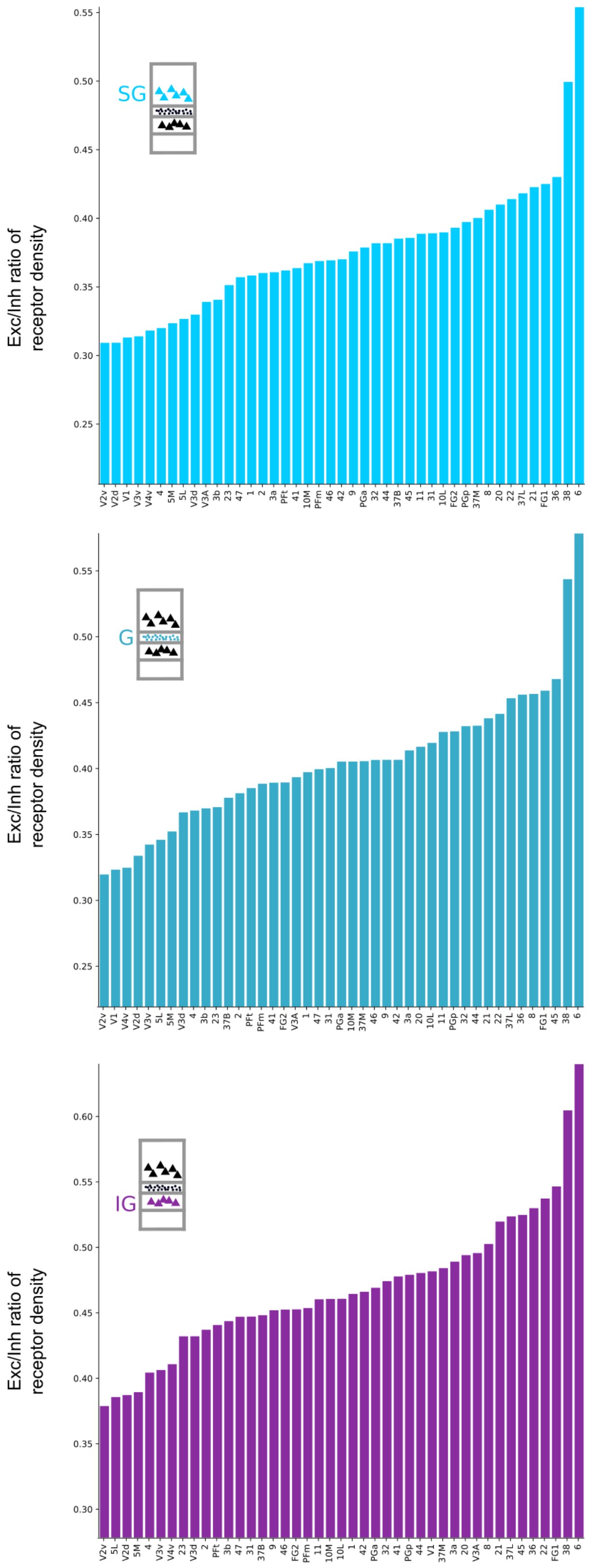


**Fig. S2.** Rank ordered areas based on their excitation/inhibition ratio in IG, G and SG layers.

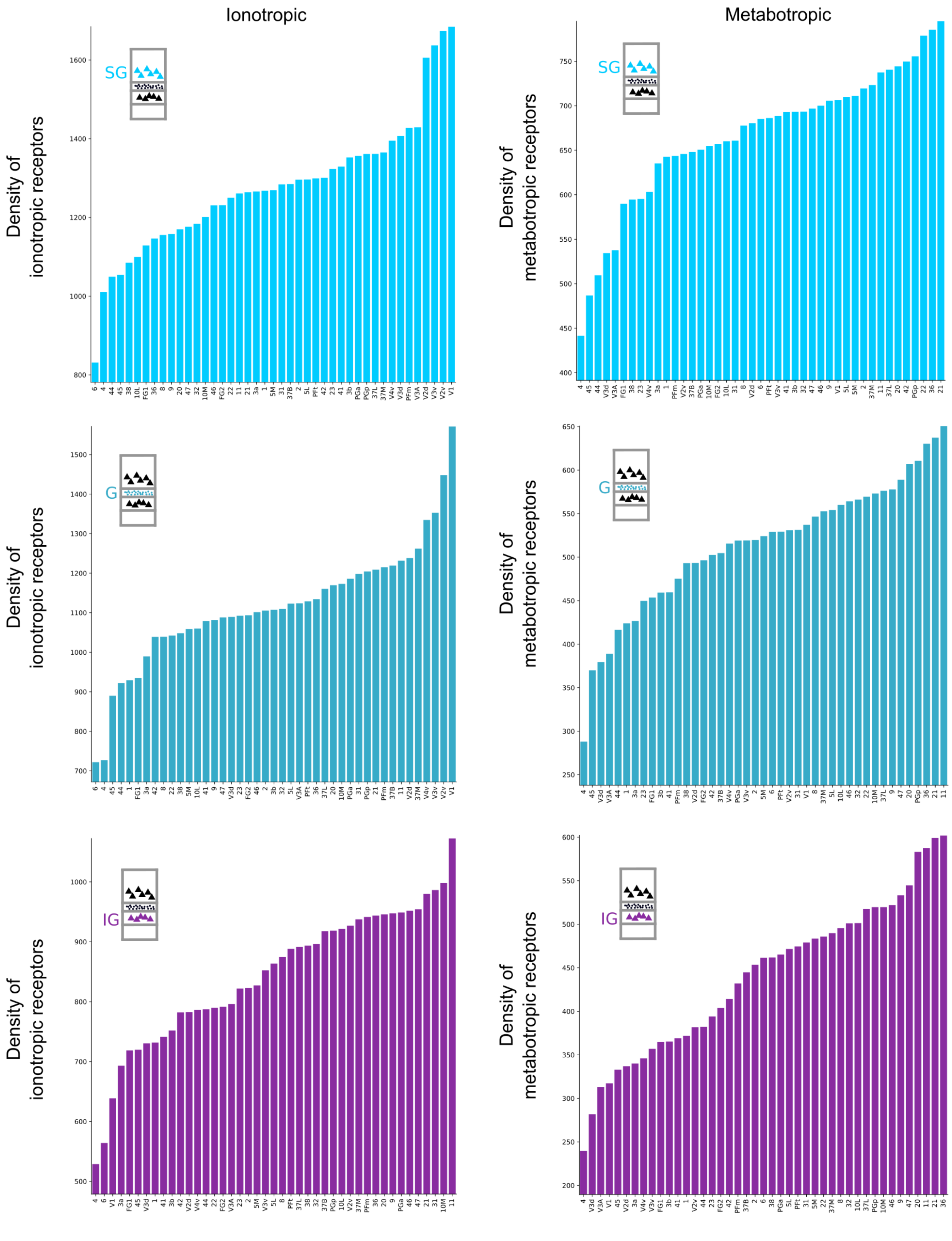


**Fig. S3.** Rank ordered areas based on the density of ionotropic and metabotropic receptors in IG, G and SG layers
